# Cellular- and systems-level profiling of amyloid-beta effects on circadian timing

**DOI:** 10.64898/2025.12.22.695794

**Authors:** Kari R. Hoyt, Tyler Kyhl, Nicklaus R. Halloy, Karl Obrietan

## Abstract

Disruption of the circadian timing system has been reported in the preclinical phase of Alzheimer’s disease (AD) and is a well-characterized component of mid- and late-stage AD. Given the distributed nature of the circadian timing system, with a central pacemaker in the suprachiasmatic nucleus (SCN) and peripheral clocks throughout the brain, understanding how AD affects this system has been challenging. To investigate how AD may disrupt circadian physiology, we focused on the amyloid-beta peptide, a key contributor to familial early-onset AD. Using the 5xFAD mouse model and *ex vivo* single-cell profiling, we examined how amyloid-beta influences clock timing in both SCN neurons and hippocampal neuronal populations. Circadian profiling of 5xFAD mice (4-and 8-months-old) showed only modest changes in key clock timing properties, including a shortening of the SCN rhythm. Interestingly, the mice showed enhanced rates of re-entrainment to changes in the light cycle, suggesting that elevated amyloid-beta levels increase the sensitivity of the SCN clock to light. Further, using both *in vitro* SCN slice explant and dispersed SCN culture models, the exogenous administration of oligomerized amyloid-beta had no significant effect on inherent clock timing capacity. In contrast, the timing properties of cultured hippocampal neurons showed a dose-dependent sensitivity to amyloid-beta. This included an elevated mesor and an increased rhythm amplitude. These findings reveal a divergence in amyloid-beta sensitivity between the central SCN clock and peripheral oscillators. This raises the possibility that circadian disruptions in AD may stem from both the destabilization and decoupling of peripheral oscillators from the central timing properties of the SCN.

## 1. Introduction

The circadian timing system is an inherent 24 hr biological oscillation that imparts a daily rhythm on nearly all physiological and behavioral processes. At the center of this timing system is an evolutionarily conserved set of autoregulatory molecular events that drive the rhythmic regulation of a limited set of ‘core-clock genes’ including the *period* (*per1-3*) and *cryptochrome* (*cry1 & 2*) gene families (Hastings et al., 2019; Mohawk et al., 2012). The circadian expression of these genes is mediated by a heterodimeric transcription factor formed by Bmal1 and Clock, which binds to the E-box motif within the 5’ regulator regions of the *per* and *cry* genes. One complete cycle of PER and CRY expression, which encompasses transcriptional activation followed by feedback repression, establishes the circadian period. This clock timing capacity occurs in a cell autonomous manner, and as such, this transcription/translation feedback loop functions independently from extracellular input (Welsh et al., 2010).

Central to clock-gated physiology is the suprachiasmatic nucleus (SCN) of the hypothalamus, which functions as the dominant pacemaker (Moore et al., 2002; Ono et al., 2024; Stephan & Zucker, 1972). The SCN provides entrainment cues in the form of synaptic and hormonal signals that set the phasing of peripheral clocks, including those found within the central nervous system. Notably, a highly coordinated cellular/systems level of clock timing, from the SCN to peripheral oscillators, is required for normal health and fitness. Consistent with this idea, recent work has revealed that the SCN works in coordination with clock timing properties within forebrain circuits to modulate cognitive processes, including learning and memory (Gerstner et al., 2009; Krishnan & Lyons, 2015; Smarr et al., 2014; Snider et al., 2018), and numerous studies have found that the disruption of this circuit, either through the loss of SCN or cortico-limbic timing, results in a marked disruption of cognitive capacity (Phan et al., 2011; Price & Obrietan, 2018; Price et al., 2016; Shimizu et al., 2016; Wardlaw et al., 2014). These data raise the possibility that the disruption of clock timing at a circuit level (e.g., peripheral oscillator fidelity and/or responsiveness to SCN output) could contribute to disorders of the nervous system where aberrant circadian-gated physiology is a comorbid feature.

Here we examined the relationship between Alzheimer’s disease (AD) pathology and the disruption of circadian timing. Work over the past several years has shown that the disruption of circadian-gated behavior and physiology (e.g., physical activity, melatonin output from the pineal gland and the sleep/wake cycle) are features of AD (Hoyt & Obrietan, 2022; Musiek et al., 2015, 2018; Wu & Swaab, 2005). To date, however, it is not clear where the locus of the circadian disruption occurs in AD; For example, is there 1) a disruption of the pacemaker properties of SCN neurons, 2) is clock output from the SCN disrupted, and/or 3) is there a disruption of timing properties within cortical and limbic circuits that underlie cognition? In this report, we explored these questions by focusing on the effects of amyloid beta (Aβ) peptide, which is thought to underlie key pathophysiological processes in AD, and other degenerative disorders of the CNS (Hampel et al., 2021; Murphy & LeVine, 2010). To this end, we used a whole-animal transgenic model (5xFAD), slice explant- and cell culture-based profiling approaches. Here we report that the SCN clock timing properties of 5xFAD mice are largely insensitive to the production and accumulation of Aβ peptides. Further, *ex vivo* profiling of 5xFAD SCN slices did not detect an effect on rhythmicity. Similarly, no effect was detected when WT SCN slices or cultured SCN neurons were treated with oligomerized Aβ peptide. In contrast, in hippocampal neurons cellular-based clock timing properties were affected by Aβ treatment. These data suggest that Aβ-induced disruptions to circadian timing in limbic and cortical circuits may contribute to the cognitive deficits seen in AD and other CNS disorders characterized by elevated Aβ levels.

## 2. Materials and methods

### 2.1 Mice

5xFAD mice (Oakley et al., 2006) were acquired from the Jackson Laboratory (MMRC #034848). To generate a breeding colony for our studies, hemizygous 5xFAD mice were crossed with WT C57BL/6J mice, and hemizygous 5xFAD and WT offspring were used. To assess cellular clock timing in the 5xFAD mouse line, hemizygous 5xFAD mice were bred with homozygous *Per1*-Venus mice (Cheng et al., 2009), which was also maintained on a C57BL/6J background. This breeding strategy generated litters that were hemizygous for the Venus transgene and hemizygous for the 5xFAD constructs. Genotyping transgenic 5xFAD and WT littermate offspring was performed by Transnetyx (Cordova, TN). All experiments involving mice adhered to Ohio State University guidelines for the ethical treatment of animals and were conducted using a protocol (Code: 2008A0227), which was approved by the Institutional Animal Care and Use Committee.

### 2.2 Circadian Activity Analysis

#### Paradigm A: Light entrainment, total darkness, and constant light assays

Wheel running activity was used to measure inherent timing and entrainment properties of the SCN. For these experiments, animals were housed individually in polycarbonate cages with running wheels (15 cm diameter), and wheel rotations were recorded in binned 5-minute intervals using the Actiview circadian analysis system (Starr Life Sciences; Oakmont, PA). Initially, animals were maintained on a 12-hour light (300 lux)/12-hour dark (LD) cycle for at least 10 days, and then a subset of the animals were challenged with a 6 hour advance and then subjected to a 6 hour delay in the lighting cycle. These same groups of animals were then maintained in constant darkness for ∼ 18 days, returned to the LD cycle, and then maintained in constant light conditions (200 lux) for ∼ 20 days. Approximately equivalent numbers of female and male mice were used for the wheel running assays, and no sex-specific differences in measurements of clock-gated activity were noted-as such, data from both sexes were pooled and used for phenotype-based analysis.

#### Paradigm B: Low light entrainment assay

Mice underwent a paradigm in which the light intensity was systematically decreased at 11-14 day intervals, and clock entrainment was assessed. To this end, mice were initially maintained on a 12-hour light (50 lux)/12-hour dark (LD) cycle, and then the light intensity was lowered to 0.5 lux, 0.10 lux, 0.05 lux and then 0.01 lux; Finally, animals were transferred to DD conditions. The Fisher’s exact test was used to compare the entrainment capacity of the two genotypes as a function of light intensity. For this analysis, the data were collapsed into two categories: Low entrainment capacity (0.5 & 0.1 lux) vs. high entrainment capacity (0.05, 0.01 lux), and a p-value of less than 0.05 was considered significant. Mice with an activity cycle that was locked to the LD cycle were considered ‘entrained’, whereas mice with a period (under LD) that was analogous to the free-running period under DD were deemed to be ‘uncoupled’-an example of this is presented in Figure 3.

### 2.3 Assessment of circadian and light-entrainment phenotypes

The quantitation of key circadian timing and clock entrainment properties (e.g., total activity, day/night activity, and free running periodicity) was performed as described previously (Cao et al., 2013; Wheaton et al., 2018) using ClockLab Analysis software (ActiMetrics; Lafayette, IN). Group means values were determined using GraphPad Prism software and presented as mean values ± SEM for each experimental group.

### 2.4 Brain Tissue Isolation

Animals were euthanized via cervical dislocation followed by rapid decapitation. For experiments that analyzed the expression of Aβ, vasoactive intestinal peptide (VIP) and arginine vasopressin (AVP), animals were sacrificed during the middle of the light cycle (zeitgeber time 6: Fig. 1); For light-pulse experiments that examined phospho-ERK expression (Fig. 4), mice were sacrificed under dim red light (< 5 lux at cage level) at circadian time 15. Brains were isolated and then cut into 500-μm-thick coronal sections using a Leica VT1200 tissue slicer (Leica; Nussloch, Germany). Tissue was then fixed with 4% formaldehyde (in phosphate buffered saline (PBS): 6 h at room temperature), followed by cryoprotection with 30% sucrose in PBS, and then cut to 44 μm with a freezing microtome. For dot-blot profiling, the SCN and cortical regions were manually dissected from freshly isolated brains that were prechilled for ∼ 1 min in oxygenated media.

**Figure 1.**
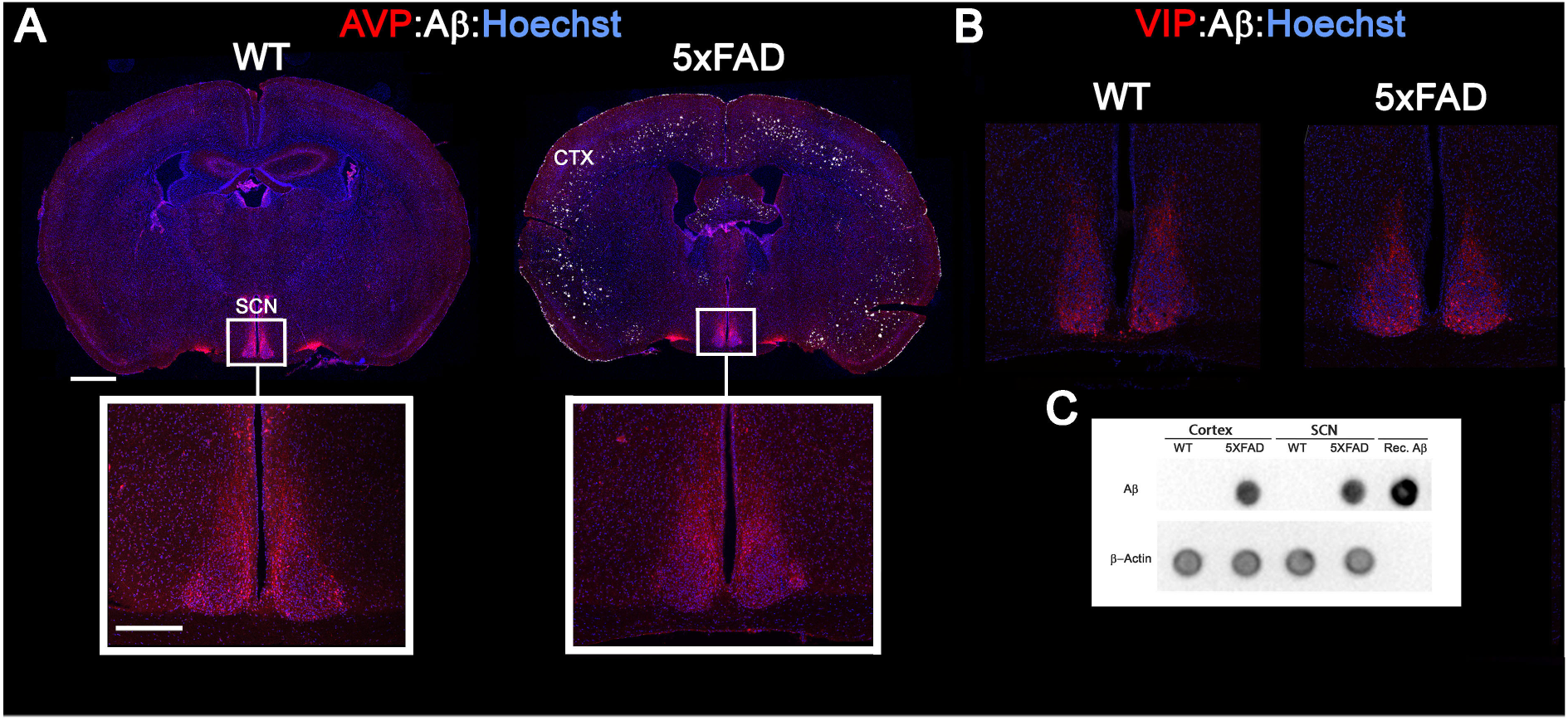
SCN morphology and Aβ expression in the 5xFAD mouse line. Coronal brain sections from 8 month-old WT and 5xFAD mice were immunolabeled for Aβ (white fluorescence) and two peptide markers of the SCN-arginine vasopressin (AVP-red fluorescence: Panel A) and vasoactive intestinal peptide (VIP-red fluorescence: Panel B)-and cellular nuclei were labeled with Hoechst (blue fluorescence). In *A*, high magnification images of the SCN are shown below each whole-brain coronal section. Representative images from 5xFAD tissue reveal marked Aβ-based plaque formation in a number of forebrain structures, including the cortex (CTX); However, plaque deposition was not observed in the SCN. Of note, AVP and VIP expression were indistinguishable between WT and 5xFAD mice, suggesting that the SCN was intact in 5xFAD animals. Bar for the low magnification image: 1000 microns; Bar for the high magnification image in A: 300 microns. C) Dot-blot protein profiling of 5xFAD and WT tissue indicates the presence of Aβ species in both the cortex and SCN of 5xFAD mice. As a control, lysates were also probed for β-actin. Further, the sensitivity of the Aβ antibody was confirmed by probing for recombinant Aβ.

### 2.5 Immunolabeling of brain sections

For fluorescence-based immunolabeling, 44 μm*-*thick tissue sections were initially permeabilized and blocked with PBS containing 0.02% Triton X-100 (v/v) and 10% bovine serum albumin (w/v) (1 hour), and then treated overnight with a rabbit anti-VIP antibody (1:2,000 dilution; Boster, Catalog code: RP1108) or a rabbit anti-AVP antibody (1:1,000 dilution: Millipore, Catalog code: AB1565). Next, sections were washed 4X in PBS and then labeled (2 hours at room temperature) with an Alexa-Fluor 488-conjugated secondary directed against the primary antibodies (1:1,000 dilution; Invitrogen). After extensive washing with PBS (6X), tissue was labeled overnight with a rabbit anti-Aβ monoclonal antibody conjugated to Alexa Fluor 594 (1:500 dilution, Catalog code: #35363). Sections were then washed 4X in PBS and labeled with the nuclei acid stain Hoechst 33342 (1:1,000 dilution, catalog code: 62249, Thermo Scientific), washed, mounted and sealed using Fluoromount-G (Thermo Fisher, Catalog code: 00-4958-02). Images of fluorescent labeling were acquired with a Leica SP8 confocal microscope (10X and 20X objectives).

For 3,3’-diaminobenzidine (DAB)-based immunolabeling, free-floating SCN tissue sections were washed in 1% PBST (PBS with Triton X-100), then treated with 0.3% hydrogen peroxide for 20 min at room temperature. After blocking with 10% normal goat serum for 1 hour, tissue was incubated overnight at 4°C with rabbit anti-pERK antibody (1:3,000; Cat #: 9,101; Cell Signaling Technology). The next day, sections were washed and incubated (2 hours) with biotin-conjugated goat anti-rabbit IgG (1:1,000; Vector Laboratories; Cat # SK-4100). Per manufacturer’s instructions, the nickel-enhanced DAB labeling kit (Vector Laboratories; Cat # SK-4100) was used to visualize the signal. Finally, sections were mounted on gelatin-coated slides, rinsed in distilled water, and sealed with Permount (Fisher Chemical). For the quantification of pERK labeling, a region of interest (ROIs) was digitally traced around the SCN core (2 sections per animal), and the intensity levels for each section were quantified using ImageJ software. SCN labeling was background subtracted using values generated from pERK labeling within the hypothalamus (∼ 500 microns lateral to the SCN). The average intensity from each animal was calculated and presented as the mean ± SEM for each genotype and condition.

### 2.6 Dot Blot Assay

SCN and cortical tissue were collected as previously described from 7–8 months old (three mice/genotype). For each genotype, pooled SCN and cortical samples were homogenized in lysis buffer (100 µL for SCN tissue: 200 µL for cortical tissue). 4-µL aliquots of each sample were applied directly onto a nitrocellulose membrane. In parallel, human recombinant Aβ (1–42) protein (Rpeptide; Catalog E1264) was prepared in lysis buffer for use as a reference. Following sample application, the membrane was allowed to air-dry and then transferred into a blocking solution consisting of 5% BSA in Tris-buffered saline with Tween 20. Membranes were subsequently incubated for 90 minutes at 4°C with either mouse anti-Aβ antibody (1:1,500; Novus Biologicals, NBP2-1307) or anti-mouse β-actin antibody (1:100,000; Phosphosolutions, 125-ACT); Membranes was washed and then exposed to an HRP-conjugated goat anti-mouse IgG secondary antibody (1:2,000; Fisher Scientific, NEF822001EA) for 2 hours at room temperature. Signal detection was performed using the Western Lightning Enhanced Chemiluminescence Substrate (PerkinElmer), and images were captured with the ChemiDoc XRS Imaging System (Bio-Rad).

### 2.7 Dissociated Neuron Preparation

Primary neuronal cultures were prepared from post-natal day 0-1 *Per1*-Venus mouse pups. For dissociated SCN cultures, brain tissue was isolated and blocked to include SCN tissue then 300 µm coronal sections were prepared using a McIlwain Tissue Chopper from which the SCN was then dissected. For dissociated hippocampal cultures, brain tissue was isolated, and the hippocampus was dissected intact from the brain. For both SCN and hippocampus preparations, after dissection, the tissue was pooled in cold dissection media (Hibernate-A (Gibco A1247501) containing 2% B-27 supplement (Gibco 17504044), 0.5 mM glutaMAX (Gibco 35050061), 100 nM MK-801 (Tocris 0924), 1% penicillin/streptomycin (Gibco 5140122) and 6 units/ml nystatin (Sigma-Aldrich N1638)). Tissue was dissociated by incubation with 2 mg/ml (∼50 units) of papain suspension (LS003126, Worthington Biochemical) in dissection media at 35°C for 30 minutes. After digestion, tissue was washed 2 times with Hibernate-A before trituration in culture media (Neurobasal-A (Gibco 1088022) containing 2% B-27 supplement (Gibco 17504044), 0.5 mM glutaMAX (Gibco 35050061), 1% penicillin/streptomycin (Gibco 5140122)) in a volume of 0.15 ml for SCN tissue and 1 ml for hippocampal tissue. SCN tissue required a more extensive trituration process than hippocampal tissue with hippocampal tissue triturated 25 times with a 1000 ul pipet tip, while SCN tissue required multiple rounds of triturations starting with 25 times with a 200 µl pipet tip, then 2 rounds of trituration ∼100 times each with a 10 µl pipet tip (with supernatant collected and pooled between triturations). After trituration and resuspension in culture media, cells were plated in 48 well culture plates (83.3923.300, Sarstedt) previously coated with poly-lysine (Gibco A38904 or Sigma-Aldrich P1274) at a volume of a 10 µl drop in the center of the well for SCN cells (density = ∼3 wells/brain) and 200 µl per well for hippocampal cells (density = ∼16 wells/brain). The cells were allowed to attach for 1 hour in a 35° C/5% CO_2_ incubator, after which the culture media was replaced with fresh culture media (350 µl/well) and the plates were returned to the incubator until long-term imaging experiments.

### 2.8 SCN Slice Preparation

SCN slice tissue was prepared from 5xFAD and WT littermates expressing *Per1*-Venus (10-11 months old) or *Per1*-Venus mice (2-4 months old) essentially as described previously (Wheaton et al., 2018). Briefly, brain tissue was isolated, blocked to include SCN, then 200-225 µm coronal sections were prepared using a McIlwain Tissue Chopper. SCN slices were transferred to fluid interface Millicell 6 well standing inserts (PICM0RG50, Millipore-Sigma, Darmstadt, Germany; 30 mm diameter transparent PTFE membrane) in 6 well plates containing culture media (Neurobasal-A (Gibco 1088022) containing 5% horse serum (Gibco 260505088), 2% B-27 supplement (Gibco 17504044), 0.5 mM glutaMAX (Gibco 35050061), 100 nM MK-801 (Tocris 0924), 1% penicillin/streptomycin (Gibco 5140122) and 6 units/ml nystatin (Sigma-Aldrich N1638)). Media were changed 1 hour after plating. Slices were then maintained in an incubator at 35°C and 5% CO_2_ until the start of long-term fluorescence imaging experiments.

### 2.9 Fluorescence Profiling of Clock Rhythms

Long-term *Per1*-Venus imaging in SCN slices and dissociated neurons was performed as previously described (Hoyt et al., 2024; Halloy et al., 2025). Culture plates were sealed with an ALA MEA-SHEET (Multi Channel Systems, Reutlingen, Germany) and transferred to a stage-top incubator (Oko Labs, Pozzuoli, Italy) mounted on an inverted Leica Stellaris confocal microscope (Leica; Nussloch, Germany) and set to 5% CO_2_ and 35°C with 95% humidity. Automated time-lapse imaging of cellular Venus transgene fluorescence (using YFP excitation/emission settings) at one-hour intervals was programmed for multiple wells per experiment. For experiments with oligomerized Aβ addition, after baseline fluorescence acquisition, slices or cells were treated with the indicated concentration of oligomerized Aβ (Human Synthetic Amyloid Beta 1-42 Oligomers: Stressmarq SPR-488, prepared according to the manufacturer’s protocol) or vehicle.

The mean Venus fluorescence intensity in SCN slices and dissociated cells was measured using Fiji/ImageJ (Schindelin et al., 2012). For SCN slices, a single ROI was drawn around the SCN and the mean fluorescence intensity was recorded for each time point. For dissociated cells, due to movement of cells with time in culture, the ImageJ plug-in, TrackMate (Ershov et al., 2022), was used to measure the mean Venus fluorescence intensity per cell. Biodare2 (Zielinski et al., 2014, 2022) was then used to detrend the time-lapse fluorescence traces and calculate the period and amplitude of Venus expression (FFT-NLLS or mFourFit method); mesor was defined as the mean Venus fluorescence of all timepoints during the noted recording time. For dissociated neurons, profiling started 6 days (SCN neurons) or 5 days (hippocampal neurons) after addition of Aβ or vehicle. GraphPad Prism was used for statistical analysis (paired and unpaired Student’s t tests as noted) and p values for statistical significance were Bonferroni corrected for multiple comparisons as noted in the figure legends.

## 3 Results

### 3.1 The 5xFAD Transgenic Mouse Line

To begin our analysis of the functional effects of Aβ on the circadian timing system we used the 5xFAD transgenic mouse line (Oakley et al., 2006). The 5xFAD transgene includes 5 mutations linked to Alzheimer’s disease: the Swedish (K670N/M671L), Florida (I716V), and London (V717I) mutations in APP, and the M146L and L286V mutations in PSEN1. In these mice, high levels of Aβ42 are detected by 1.5 months of age, and amyloid plaques and gliosis are detected at two months of age. Further, synaptic degeneration and neuronal loss are detected by four to six months of age, respectively (Eimer & Vassar, 2013; Oakley et al., 2006). These mice also exhibit a spectrum of cognitive and motor impairments (Forner et al., 2021; Girard et al., 2013; Jawhar et al., 2012), and limited work to date has examined SCN clock timing properties (King et al., 2025; Song et al., 2015).

Initially, immunofluorescence-based profiling was used to examine Aβ plaque deposition in the CNS of 8 month-old 5xFAD and WT mice. Representative images (Fig. 1A), reveal marked Aβ-based plaque formation in a number of forebrain structures, including the cortex and striatum, however, limited plaque deposition was observed in the hypothalamus, including the SCN. Further, with respect to the SCN, tissue was immunolabeled for two peptidergic markers: arginine vasopressin (AVP), which labels dorsal and lateral cell populations (referred to as the ‘SCN shell’) and vasoactive intestinal peptide (VIP), which labels a distinct ventral cell population (referred to as the ‘SCN core’). Notably, AVP and VIP expression (Fig. 1A and 1B, respectively) within the SCN did not appear to be affected in 5xFAD mice; Hence, marked expression of both peptides were observed, and the pattern of expression within the SCN was indistinguishable between 5xFAD and WT mice. In addition, we examined Aβ expression using dot-blot profiling (Fig. 1C). To this end, SCN and cortical tissue were probed using an antibody against the oligomerized form of Aβ. Representative data reveal that in both the SCN and cortex, marked Aβ was detected in the 5xFAD mouse line, but not in WT animals. Further, the sensitivity of the assay was confirmed by probing for expression of recombinant Aβ. Together, these data indicate that the SCN is morphologically intact, and that oligomerized Aβ is expressed in SCN of 5xFAD mice.

### 3.2 5xFAD: Circadian Profiling

To assess the potential effects of Aβ on the circadian timing system, we first examined the timing properties of the SCN in 5xFAD mice. Wheel running activity was used as a functional output of the SCN clock, and analysis was performed at 4 months- and 8 months-of-age; a time frame that denotes a marked acceleration of the pathological process (Oakley et al., 2006). Mice were initially placed on a standard 12 hour light/12 hour dark cycle, and the entrainment properties of 5xFAD and WT mice were profiled over a 10-day period (Fig. 2A and 2B). As expected for WT mice, the majority of locomotor activity occurred during the night-time domain. 5xFAD mice exhibited a similar locomotor activity profile, with the vast majority of wheel running activity consolidated to the dark phase. A comparison of 5xFAD and WT mice at 4 months- and 8 months-of-age did not detect a significant difference in the amount of daily activity nor differences in the amount of activity during the light and dark periods (Fig. 2C and 2D; Table 1).

**Figure 2.**
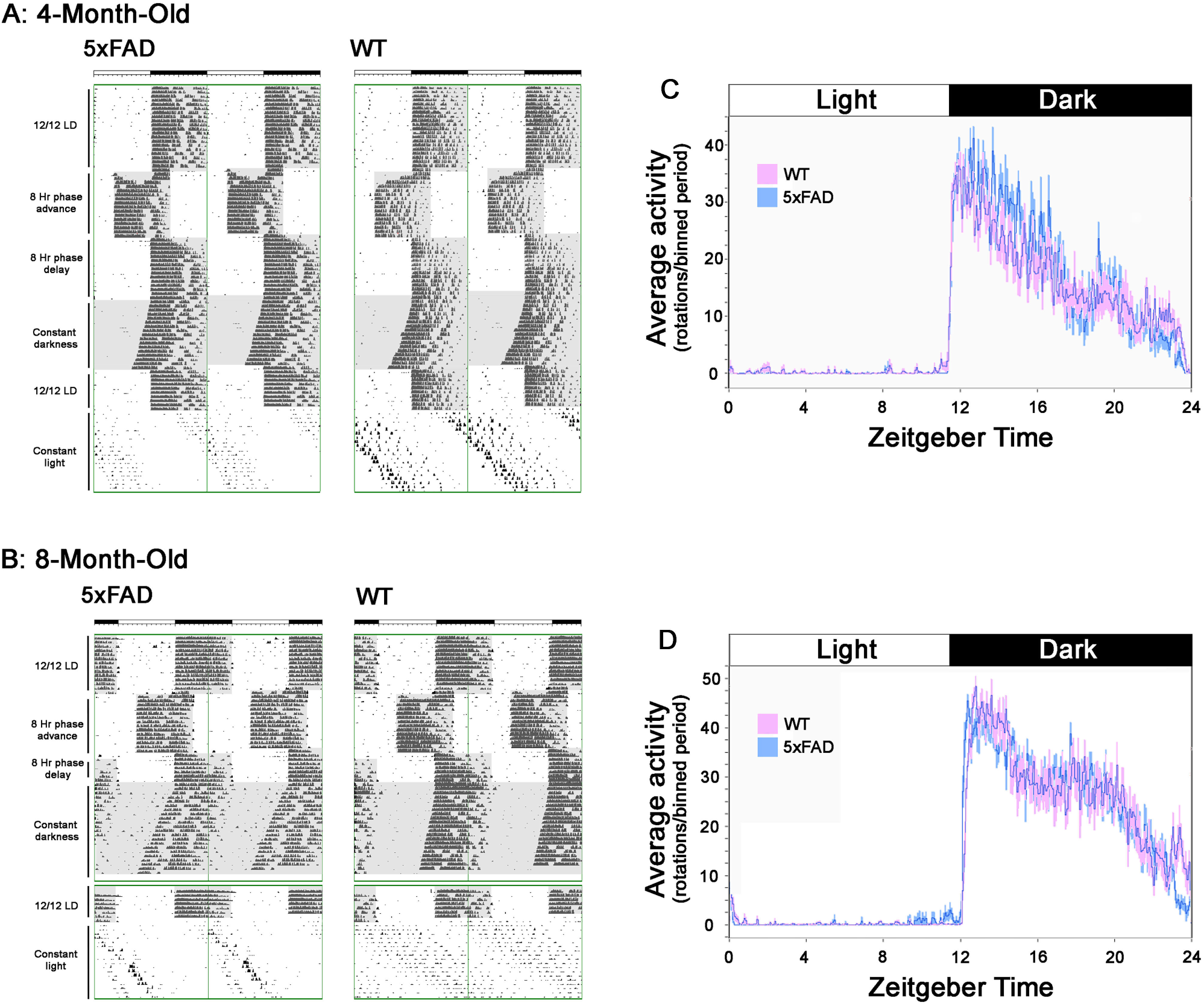
SCN-based circadian timing is largely unaffected in the 5xFAD transgenic mouse line. A & B: Representative double-plotted actographs of wheel-running activity in 5xFAD and WT littermate mice. Analysis was initiated at 4 months-of-age (*A*: early CNS pathogenic period) and at 8 months of age (*B*: marked CNS pathogenesis). Initially, mice were entrained on a 12 hour LD cycle (400 lux; denoted as the “12/12 LD”) and then tested for light-evoked clock resetting (using a ‘jet lag’ reentrainment paradigm), where animals were transferred to an 8 hour phase-advance LD cycle, and then transferred back (phase-delay) to the original LD cycle. After stable reentrainment, the inherent pacemaker activity of the SCN was profiled by transferring mice to constant darkness for ∼ 20 days (denoted as the ‘Constant darkness’). Mice were then returned to a standard LD cycle, and once stable entrainment was achieved, the robustness and fidelity of the SCN timing system was tested by maintaining animals under constant light (LL: 200 lux). Gray areas indicate the periods when the light was off and white areas indicate periods when the light was on. Of note, the 2 separate graphs for the 8-month-old traces (B) were separated by an ∼ 2 week period when the activity was not recorded. (C & D) Mean daily locomotor activity profiles of 5xFAD (n=8/age group) and WT (m=8/age group) littermate mice in LD. Of note, at both ages (4-months and 8-months-of-age), both the total amount of activity and the day/night distribution of activity are similar for 5xFAD and WT mice.

**Table 1:** Circadian and clock entrainment properties of 5xFAD and WT mice (representative wheel-running activity are presented in *figure 2*). * = p < 0.05; ** = p < 0.01, Student’s t tests.

| Parameter (mean ± standard error of the mean) | Wild-Type (n=8) | 4-month old<br>5XFAD (n=8) | p value | Wild-Type (n=8) | 8-month old<br>5XFAD (n=8) | p value |
| --- | --- | --- | --- | --- | --- | --- |
| 1. Overall activity in LD (rotations/5 min) | 48.87 ± 7.44 | 56.77 ± 7.19 | 0.439 | 72.14 ± 10.00 | 71.23 ± 6.55 | 0.940 |
| 2. Average acrophase in LD | 4.16 ± 0.30 | 3.89 ± 0.11 | 0.417 | 4.44 ± 0.12 | 4.28 ± 0.30 | 0.652 |
| 3. Activity during the light (L) period (rotations/5 min) | 3.68 ± 1.57 | 1.37 ± 0.36 | 0.204 | 1.94 ± 0.41 | 2.16 ± 1.28 | 0.869 |
| 4. Activity during the dark (D) period (rotations/5 min) | 63.61 ± 8.55 | 76.34 ± 8.03 | 0.302 | 143.58 ± 20.34 | 140.61 ± 13.20 | 0.904 |
| 5. Light/dark activity ratio | 0.04 ± 0.01 | 0.01 ± 0.00 | 0.101 | 0.02 ± 0.01 | 0.02 ± 0.01 | 0.783 |
| 6. Variance in activity onset in LD (min) | 7.13 ± 1.06 | 5.57 ± 0.69 | 0.255 | 7.57 ± 1.73 | 5.50 ± 1.32 | 0.352 |
| 7. Days to re-entrain to 8h advancing LD cycle | 8.00 ± 1.07 | 5.00 ± 0.57 | 0.027* | 5.13 ± 1.33 | 4.14 ± 0.67 | 0.524 |
| 8. Days to re-entrain to 8h delaying LD cycle | 3.13 ± 0.40 | 1.75 ± 0.25 | 0.013* | 1.88 ± 0.48 | 1.14 ± 0.14 | 0.181 |
| 9. Circadian period ($\tau$) in DD (hr) | 23.77 ± 0.06 | 23.71 ± 0.04 | 0.450 | 23.75 ± 0.05 | 23.61 ± 0.05 | 0.047* |
| 10. Overall activity in DD (rotations/5 min) | 43.25 ± 8.68 | 59.14 ± 9.33 | 0.232 | 83.31 ± 13.18 | 47.55 ± 8.30 | 0.038* |
| 11. Alpha duration in DD (hrs) | 10.13 ± 0.42 | 9.31 ± 0.76 | 0.361 | 10.54 ± 0.47 | 10.95 ± 0.58 | 0.595 |
| 12. Rho duration in DD (hrs) | 13.87 ± 0.42 | 14.69 ± 2.02 | 0.361 | 13.46 ± 0.47 | 13.05 ± 0.58 | 0.595 |
| 13. Acrophase (activity peak) during DD | 4.55 ± 0.35 | 4.47 ± 0.33 | 0.878 | 4.87 ± 0.35 | 5.29 ± 0.29 | 0.409 |
| 14. Circadian period ($\tau$) in LL (hr) | 24.86 ± 0.06 | 24.73 ± 0.06 | 0.006** | 25.34 ± 0.06 | 25.29 ± 0.09 | 0.677 |
| 15. Overall activity in LL (rotations/5 min) | 1.20 ± 0.43 | 5.72 ± 1.40 | 0.161 | 13.39 ± 6.88 | 2.03 ± 0.72 | 0.144 |

To assess whether Aβ affects the inherent timekeeping properties of the SCN, mice were transferred to total darkness (DD) and allowed to ‘free-run’. Under this condition, the period of wheel running behavior is set by the SCN oscillator, and as such, this rhythm can be used to measure the inherent periodicity of the SCN clock. At both 4 months- and 8 months-of-age, 5xFAD mice exhibited a slightly shorter free running period, which reached statistical significance (p < 0.05; ∼ 8 min difference) at the 8 month-old time point (Table 1). Other parameters, including total activity levels, and the timing of acrophase were not significantly different between the two genotypes for each age in DD.

The timekeeping properties of the SCN were also examined using a constant light (LL) paradigm. LL has been shown to disrupt clock timing at a systems-level within the SCN, and thus lead to a number of marked changes in SCN-gated locomotor activity, including the lengthening of the period, a reduction in locomotor output, and fragmentation of the running rhythm (Daan & Pittendrigh CS, 1976; Pittendrigh, 1960). Using a 200 lux constant light paradigm, WT mice exhibited the expected decrease in overall locomotor activity (relative to DD conditions) and a lengthening of the circadian period (Fig. 2A and 2B). Similar to this, 5xFAD mice had reduced overall activity and a lengthening of the circadian period; Interestingly, however, the LL-induced tau lengthening in 5xFAD mice was significantly shorter than in WT mice at 4 months-of-age, and a similar trending pattern was also observed at 8 months-of-age (Table 1).

### 3.3 5xFAD: Light-mediated clock entrainment

The phasing of the SCN clock is tightly controlled by the 24-hour light cycle; As such, any alterations in the sensitivity of the clock to light could have profound effects on clock-gated physiology and behavior. Notably, individuals with dementia have been reported to show an increased sensitivity to light (Barrick et al., 2010), thus raising the possibility that AD patients have altered/heightened sensitivity of the SCN clock to light. To examine this question in the 5xFAD mouse line, we performed a series of light entrainment experiments, where mice were subjected to an 8 hour phase-advance and an 8 hour phase-delay of the lighting cycle and the rate of reentrainment was determined (Fig 2A and 2B). Interestingly, in the 4 month-old mice, the 5xFAD line entrained at a significantly faster rate than WT mice to the 8-hour advance in the LD cycle (Table 1). Eight month-old mice also showed a faster rate of reentrainment than WT mice, although these data did not reach statistical significance. Similarly, in 4 month-old mice, 5xFAD mice reentrained to an 8 hour delay significantly faster than WT mice, and 8 month-old mice showed a similar pattern, but the difference in the rate of reentrainment for the two genotypes did not reach statistical significance (Table 1). Together, these data suggest that 5xFAD mice show a heightened relative level of sensitivity to the entraining effects of light.

To further examine light sensitivity, we performed a series of experiments to assess the lower limits of the entraining capacity. To this end, 10 month-old 5xFAD and WT mice were tested for entrainment capacity to a series of step-like reductions in light level-from 50 lux to 0.01 lux-over 10 day intervals (Fig. 3A). Interestingly, on average, the 5xFAD mice exhibited entrainment capacity at much lower light levels than WT mice (Fig. 3A and 3B). Along these lines, 5 of 8 5xFAD mice were able to maintain entrainment down to 0.01 lux of light, whereas only 3 of 8 WT mice maintained entrainment. However, a significant difference in the sensitivity to light was not detected using Fisher’s exact test (Fisher’s exact p = 0.12), although, the odds ratio (11.67), suggests a trend toward 5xFAD mice showing significantly enhanced light entrainment capacity. These data, along with the jet lag entrainment data, raise the prospect that 5xFAD mice exhibit heightened SCN entrainment capacity.

**Figure 3:**
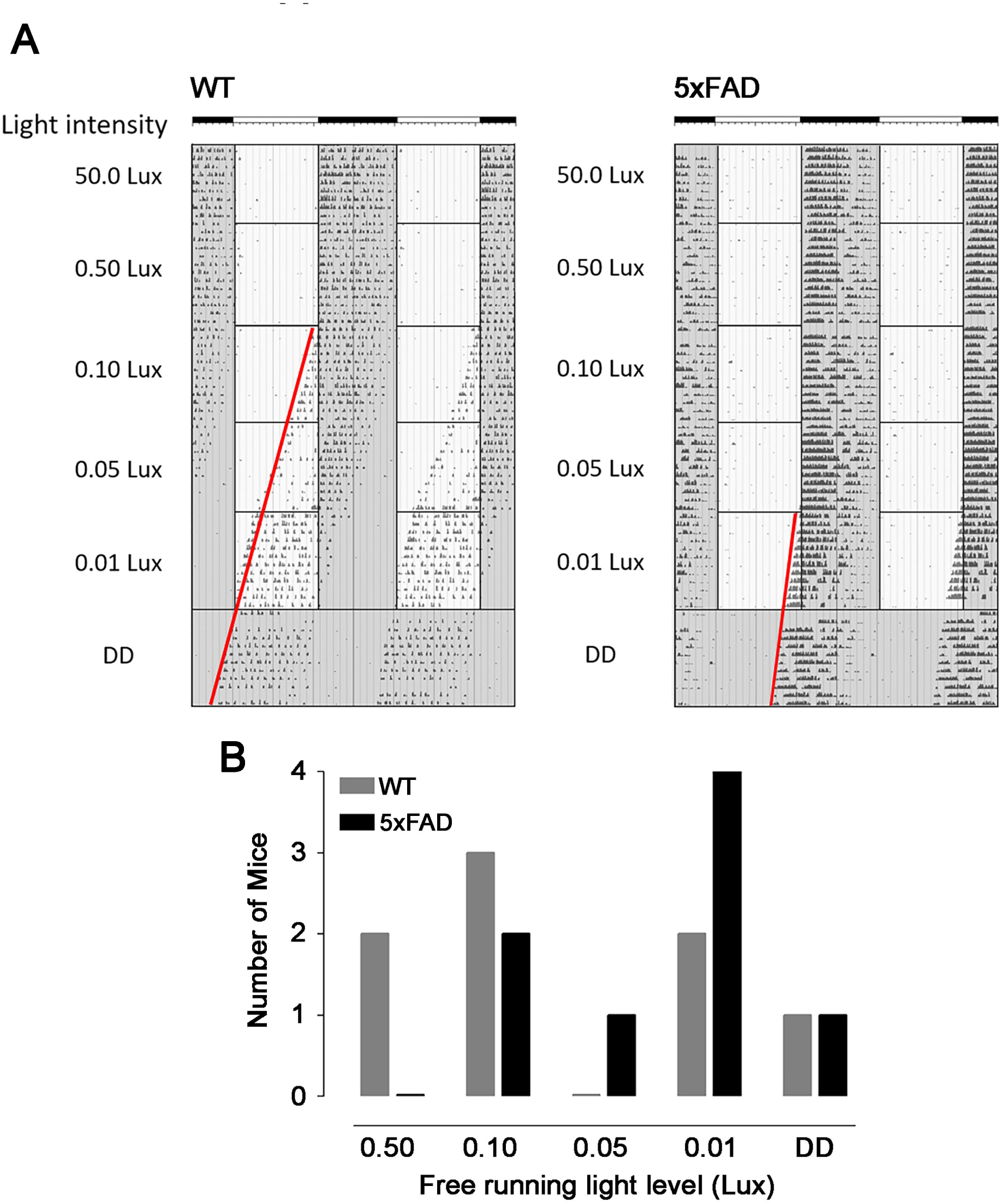
Light-sensitivity in 5xFAD mice. A: SCN entrainment capacity of 5xFAD and WT littermate mice was tested by exposing animals to a step-wise reduction in light levels over multi-day periods. Gray areas indicate the periods when the light was off and white areas indicate periods when the light was on. Representative actographs show a 5xFAD mouse that maintains SCN clock entrainment down to 0.05 lux, whereas the WT mouse was only able to maintain entrainment down to 0.50 lux. Loss of entrainment (i.e., free running) was revealed when the onset of locomotor activity began during the illuminated period of the 12/12 LD cycle. As confirmation of the loss of light entrainment, animals were transferred to DD at the end of the experiment, and the free-running period (tau) was assessed. Regression analysis (denoted by the red lines) reveals that mice did not use light as a zeitgeber at the noted light levels. B: Entrainment capacity as a function of the light levels in WT (n=8) and 5xFAD (n=8) mice. The Y-axis denotes the light level at which animals lost entrainment capacity and initiated *free running*. For the *DD* group, mice were able to maintain entrainment down to the lowest light level (0.01 lux), and only free ran under total darkness (DD).

### 3.4 5xFAD: Light-mediated MAPK pathway activation

To examine possible mechanisms that underlie the enhanced sensitivity of the 5xFAD line to light, we profiled the photic response properties of the SCN at a cellular level. To this end, we examined light-evoked activation of the MAPK signaling cascade in the SCN. The MAPK cascade is a critical photic input pathway that underlies the resetting capacity of the SCN clock (Butcher et al., 2002; Obrietan et al., 1998). For these experiments mice were dark-adapted for two days and then exposed to light (10 lux, 10 minutes) at CT17. Animals were sacrificed following the termination of the light stimulation, and SCN containing tissue was profiled via immunohistochemical labeling for the phospho-activated form of ERK (a marker of MAPK cascade activity). Representative images reveal light-evoked ERK activation in both 5xFAD and WT mice (Fig. 4A). Quantitative densitometric analysis of the SCN core did not detect a significant difference in the level of light-evoked ERK activation between the two genotypes. Further, baseline ERK activation was not different between the two genotypes (Fig. 4B). Together, these data suggest that the SCN neuronal response to light is not markedly affected in 5xFAD mice.

**Figure 4:**
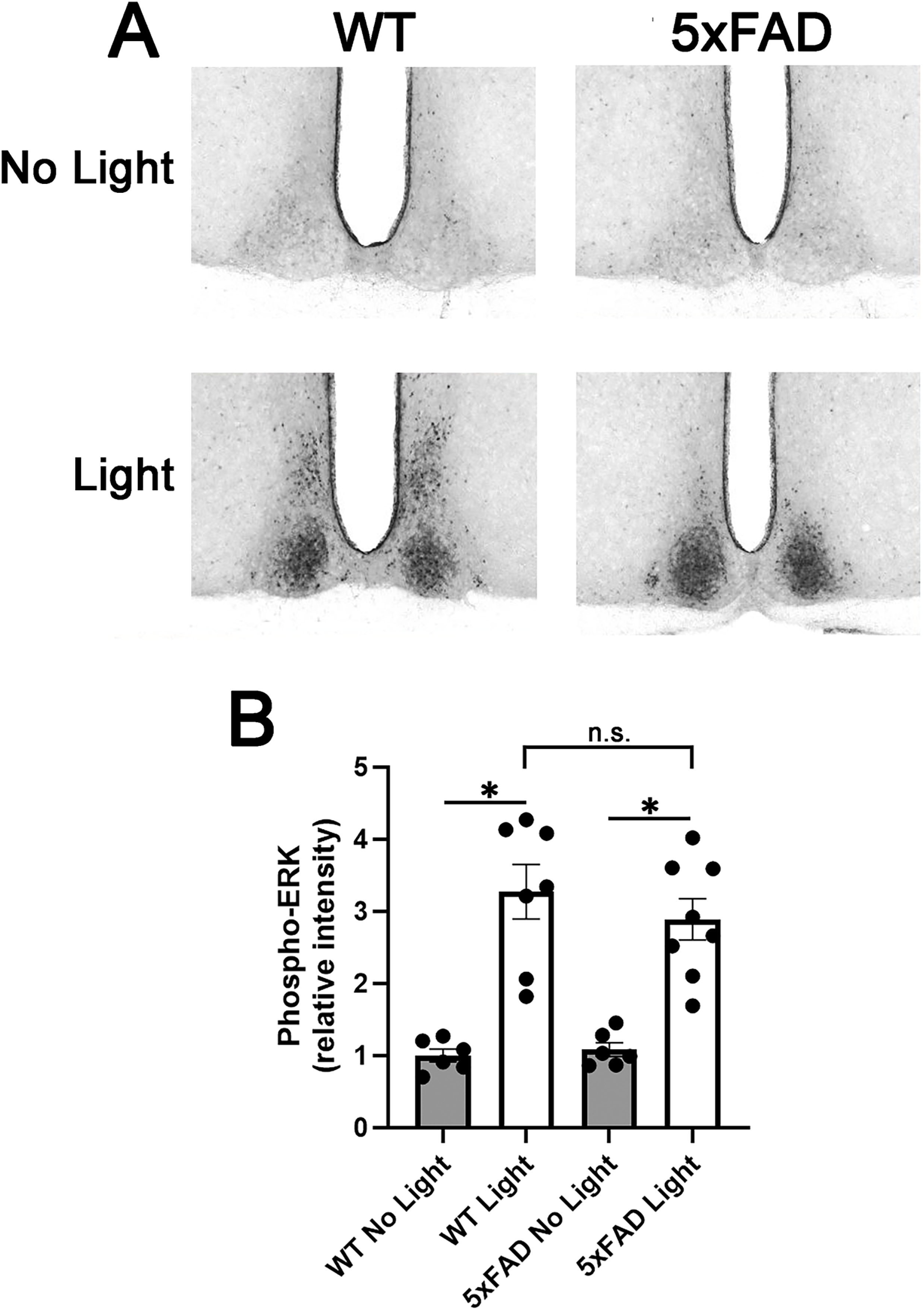
Light-evoked ERK activation in the SCN is not altered in 5xFAD mice. A: Representative immunohistochemical labeling for phospho-ERK (pERK) in 5xFAD and WT mice. Animals were dark-adapted for 2 days and then either sacrificed (control animals “No Light”) or exposed to light (10 lux; “Light”) for 5 min at CT17. Relative to control animals, light triggered marked light-evoked pERK in the SCN of both 5xFAD and WT animals. B: pERK quantitation for the control and light treatment conditions. The pERK expression in the WT SCN under “no light” condition was set to a value of ‘1’ and expression levels in all other conditions are reported as relative values. * = p < 0.01, significantly different from the corresponding *No Light* condition, ANOVA with Bonferroni’s Multiple Comparisons test. n.s.: significant differences in evoked pERK levels were not observed between the WT and 5xFAD animals. Error bars denote the SEM. Each data point represents the pERK value from a single animal; Data were averaged from 6-7 animals per condition.

### 3.5 : *In vitro* profiling of SCN timing

To provide a more mechanistic dissection of the potential effects of Aβ on the functional properties of the SCN timing system, we pursued a combination of organotypic slice and dispersed neuronal cell culture-based profiling approaches. These experimental approaches employed our *Per1*-Venus transgenic reporter mouse line (Cheng et al., 2009; Hoyt et al., 2024). The *Per1*-Venus transgene provides a dynamic, fluorescence-based read-out of transcriptional activity from the *period1* promotor. Given that the rhythmic expression of Period1 is a critical component of the core clock TTFL, cellular-based transcriptional monitoring from the *period1* locus serves as an effective readout of the circadian timing system. Initially, for these experiments, mice transgenic for the *Per1*-Venus construct were crossed to the 5xFAD mouse line and organotypic SCN slices were prepared from 10- to 11-month-old-mice and profiled over a 5-day period (Fig. 5A-C). Confocal-based Venus profiling for the circadian period (tau), and rhythm amplitude did not detect a significant effect of the 5xFAD transgene, relative to WT rhythms (Fig. 5D and 5E). To further our examination of the potential effects of Aβ on SCN timing, we performed a series of experiments where the oligomerized form of Aβ was directly applied to *Per1*-Venus SCN explant tissue. To this end, tissue from *Per1*-Venus mice were prepared from 2- to -4 month-old animals, treated with 4 μM oligomerized Aβ (or vehicle), and then profiled for 4 days (Fig. 5F). Interestingly, similar to the data from the 5xFAD mouse line, oligomerized Aβ did not affect SCN clock periodicity or amplitude (Fig. 5G and 5H) relative to vehicle-treated slices. Of note, a decrease in the amplitude of the rhythms as a function of the time in culture was observed for both the vehicle- and Aβ-treated; Importantly, relative to vehicle-treatment, Aβ treatment did not affect the magnitude of this decrease in rhythm amplitude. Together, these data indicate that key rhythm-generating properties of the SCN are not markedly affected by Aβ.

**Figure 5:**
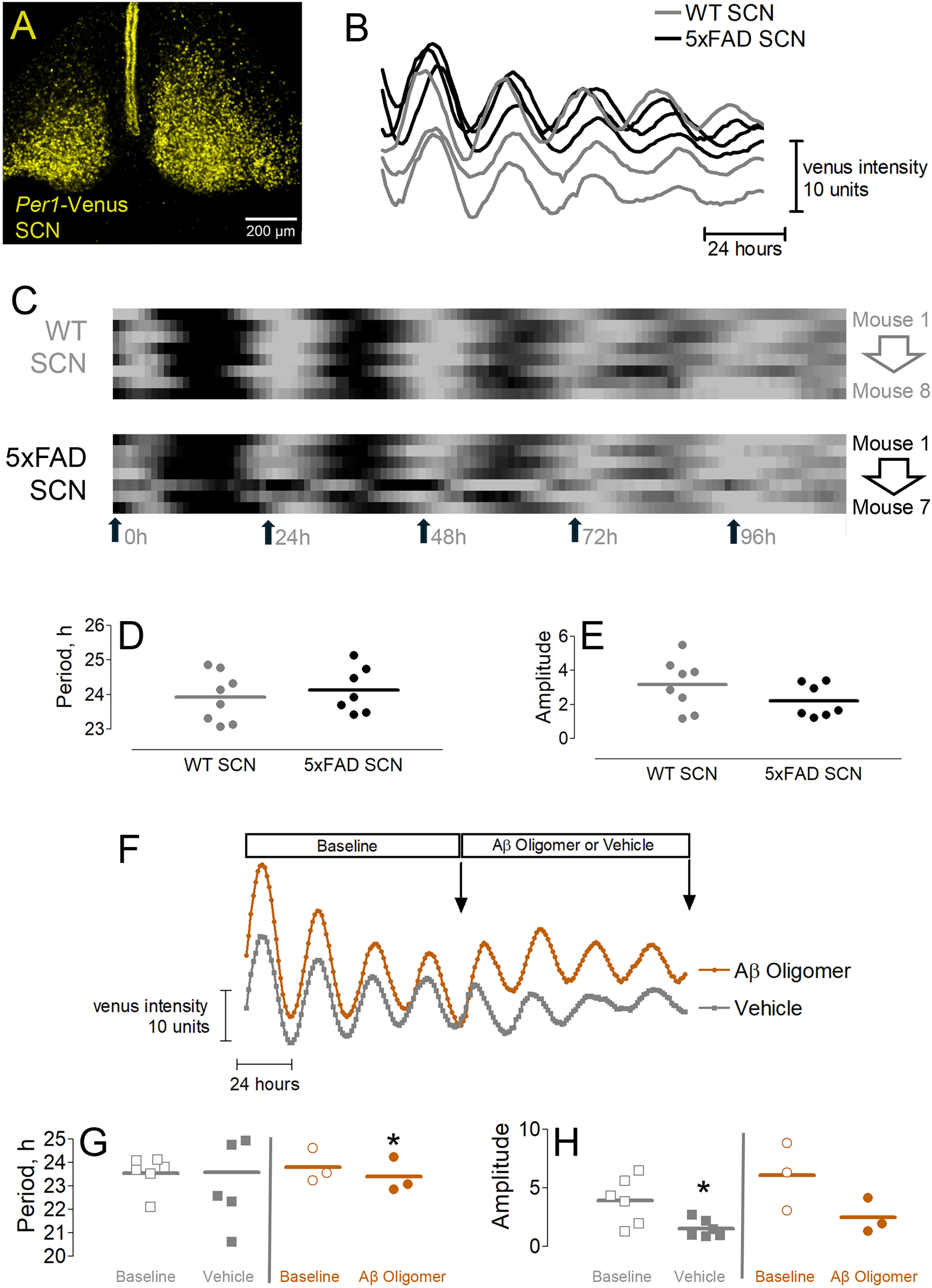
Aß and the *in vitro* profiling of SCN timing. A: Representative fluorescence image of *period1*-Venus reporter expression from an SCN slice explant. B: Detrended SCN Venus expression profile for 5xFAD mice (n = 3) and non-transgenic (WT) littermates (n = 3) (10-11 months-old). Data were collected at 1 hour intervals. C: Raster plot profiling of Venus expression in 5xFAD mice (n = 7) and WT littermates (n = 8). High Venus expression (top 90^th^ percentile) is denoted by black color, middle 50^th^ percentile is medium gray, and the lowest 10^th^ percentile is light gray. Venus rhythm period (D) and amplitude (E) were quantified. Of note, no significant differences between 5xFAD mice and WT littermates were observed (Student’s t test). Each point represents a single SCN explant. F: Representative profiling data from *Per1*-Venus slices treated with Aß oligomers (4 µM) or vehicle. Representative traces are from one SCN explant for each condition. Relative to baseline profiling and vehicle administration, treatment with Aß did not affect rhythm periodicity (G) or rhythm amplitude (H): n = 3 for Aß treatment and n = 6 for vehicle treatment. For each SCN explant, paired t tests were performed to assess the effect of treatment (vehicle or Aß treatment) compared to baseline (G) and (H); * denotes a significant difference from the corresponding baseline value. To assess whether Aß treatment was different from vehicle treatment, the treated values was subtracted from the baseline values for each SCN explant, and then this group mean difference was compared by Student’s t test for vehicle and Aß treatment conditions; No statistically significant difference was detected when comparing vehicle to Aß treatment for (G) nor (H). Significance level was set to p<0.017 using a Bonferroni correction for multiple comparisons.

Notably the slice-based profiling approaches outlined above provide a read-out of circuit level SCN timing, which is dependent on a combination of cellular clocks, and synaptic and paracrine signaling. To further our investigation of the potential effects of Aβ on SCN timing, we eliminated the synaptic and paracrine component of the timing system and focused specifically on the autonomous oscillatory properties of SCN neurons. To this end, dispersed cultures of SCN neurons were prepared from the SCN of *Per1*-Venus pups (P0-1). After 6 days in culture, tetrodotoxin (1 μM) was added to the media (to suppress action-potential-mediated transmitter release) and cellular-level clock timing was profiled at 1-hr intervals over a 6 day period. As expected, SCN neurons exhibited robust rhythm generating capacity (Fig. 6A and 6B). Following this baseline profiling period, cells were treated with oligomerized Aβ (1 μM or 4 μM) or vehicle (0 μM) and single-cell Venus rhythms were profiled. Interestingly, oligomerized Aβ did not affect the periodicity, amplitude, or rhythm mesor of cellular oscillators (Fig. 6C-6E). Importantly, as a function of time in culture, an increase in the mesor of the Venus rhythm was observed (Fig. 6D); However, relative to vehicle-treatment, Aβ did not significantly alter the magnitude of this incremental increase. Collectively, these data indicate that clock timing in the SCN is largely resistant to the effects of Aβ, and as such, raise the prospect that Aβ could affect clock-gated physiology via the disruption/dysregulation of ancillary (extra-SCN) CNS oscillator populations.

**Figure 6:**
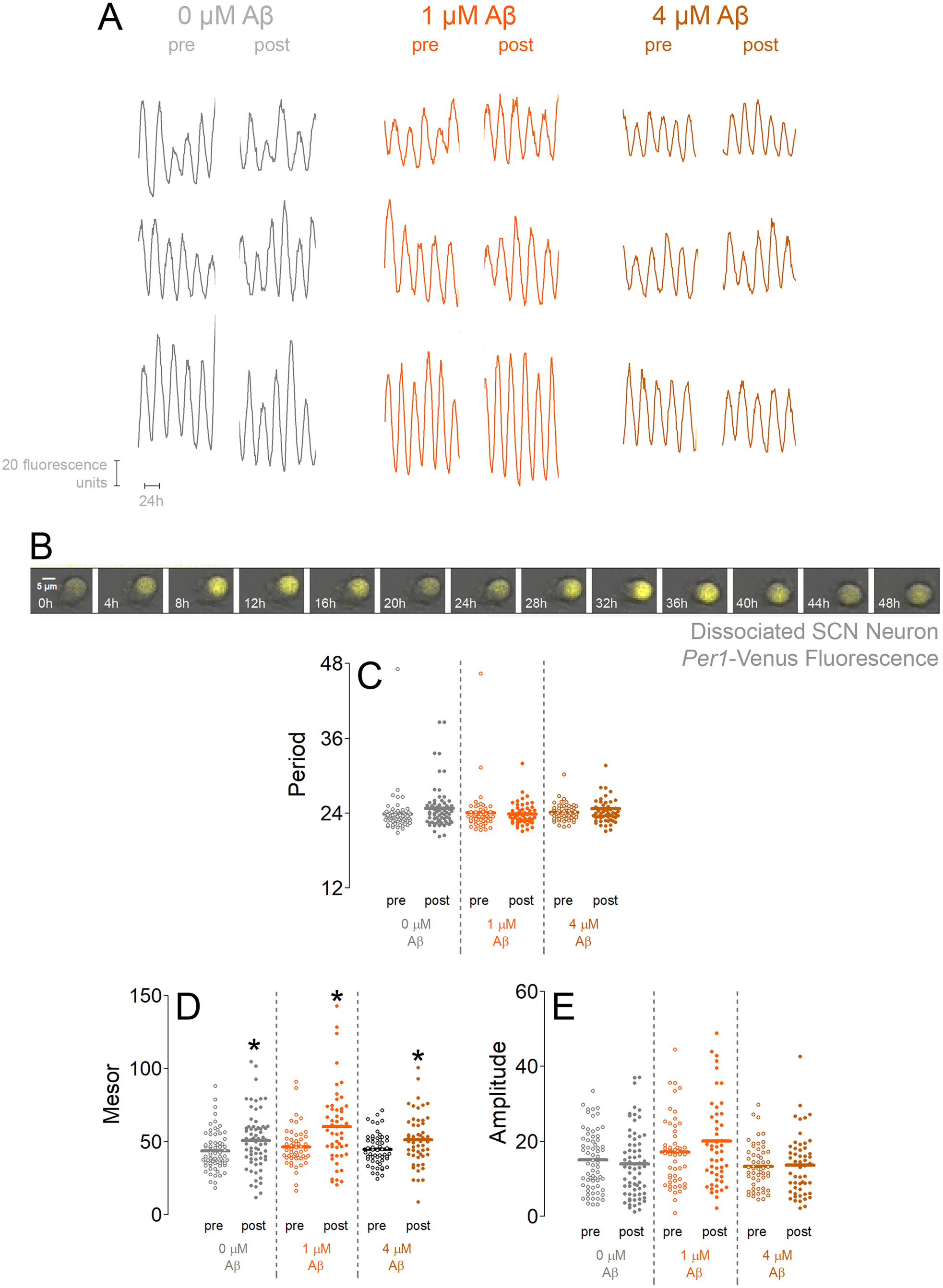
Administration of oligomerized Aß does not alter the timekeeping properties of cultured SCN neurons. SCN neurons were isolated, plated and maintained in TTX-containing media. A: Representative Venus fluorescence traces (n = 3 per treatment group) of isolated SCN neurons before (“pre”) and during (“post”) Aß treatment with either vehicle (0 µM) or oligomerized Aß (1 µM or 4 µM). B: Representative time-lapse profiling of Venus expression for a single SCN neuron. Venus expression (yellow (YFP) fluorescence channel) was merged with the corresponding brightfield image. Of note, Venus expression showed marked rhythmicity, with a periodicity of ∼ 24 hours (the corresponding Venus fluorescence trace for this representative SCN neuron is the bottom “pre” trace for the 0 µM Aß treatment in “A”. Period (in hours: C) and amplitude (E) were similar for each of the 0, 1 and 4 µM Aß conditions both before (“pre”) and after (“post”) treatment, while mesor (D) significantly increased with time in culture for all treatment conditions. For statistical analysis, paired t tests were used to compare baseline (“pre”) and treated (“post”) measurements for each SCN neuron to determine the group mean effect of treatment (vehicle: 0 µM; Aß: 1 µM or 4 µM) for panels C, D, and E. * denotes a significant difference between the corresponding “pre” and “post” values. To assess differences across the 0 µM, 1 µM or 4 µM treatments groups, the “post” value was subtracted from the “pre” value for each group, and then these differences were compared using the Student’s t test (0 µM vs. 1 µM or 4 µM; and 1 µM vs. 4 µM); No statistically significant differences were detected for these comparisons. Significance level was set to p<0.0083 using a Bonferroni correction for multiple comparisons. Each datapoint represents a single SCN neuron; n = 67 neurons for 0 µM, 51 neurons for 1 µM, and 54 neurons for 4 µM Aß.

### 3.6: Aβ and the *in vitro* profiling of hippocampal neurons

As noted, clock timing is a distributed process, where peripheral oscillators work in conjunction with the SCN to shape physiology and behavior. Here, we examined whether clock timing within forebrain neurons that underlie learning, memory and cognition could be affected by oligomerized Aβ. To this end, we utilized the dispersed cell culture model described above, where hippocampal neurons were isolated from *Per1-*Venus pups (P0-P1), treated with TTX (1 μM), and then profiled for the effects of oligomerized Aβ (1 μM and 4 μM) on cell-autonomous clock timing capacity. For these assays, we first recorded baseline rhythmicity for five days (the pretreatment condition). Cultures were then exposed to oligomerized Aβ for ten days, and rhythmicity was assessed during the final five days of this treatment period.

Initially, time-lapse profiling of Venus expression revealed that cultured hippocampal neurons exhibited a range of oscillatory capacity (from relatively consistent, albeit weak, rhythms, to rhythms that were erratic) with respect to both the periodicity and consistency (Fig. 7A and 7B). Further a subset of the neurons exhibited long oscillations (> 36 hours), which is well-beyond the expected circadian timeframe of dispersed cell-autonomous oscillations that have been reported for the SCN (e.g., 20-30.9 hrs: Honma et al., 2004; 21.25- 26.25 hr: Welsh et al., 1995). Using a periodicity range from 16-50 hrs, 83% of hippocampal neurons (n=334) were found to be rhythmic. Further, cellular-level longitudinal profiling revealed a consistent lengthening of the circadian period over the duration of the profiling (e.g., Fig. 7A and 7C: 0 μM Aβ (vehicle) condition). Here, it is worth noting that the weak inherent oscillatory activity of cultured hippocampal neurons may, in part, reflect their role as ancillary oscillators that require input from the SCN (e.g., a daily entrainment cue); Accordingly, disruption of clock timing in the SCN abrogates clock rhythms in forebrain circuits (Rath et al., 2013). We found that the exogenous application of oligomerized Aβ (1 μM and 4 μM) did not affect periodicity (Fig. 7C). Interestingly, however, 4 μM Aβ treatment triggered a significant increase in rhythm mesor and amplitude, relative to the vehicle (0 μM) and 1 μM treatment conditions (Fig. 7D and 7E). Together, these results suggest that hippocampal neurons, but not SCN neurons, exhibit altered clock timing properties in response to Aβ treatment, indicating a potential site of circadian vulnerability in Alzheimer’s disease and related Aβ pathologies.

**Figure 7:**
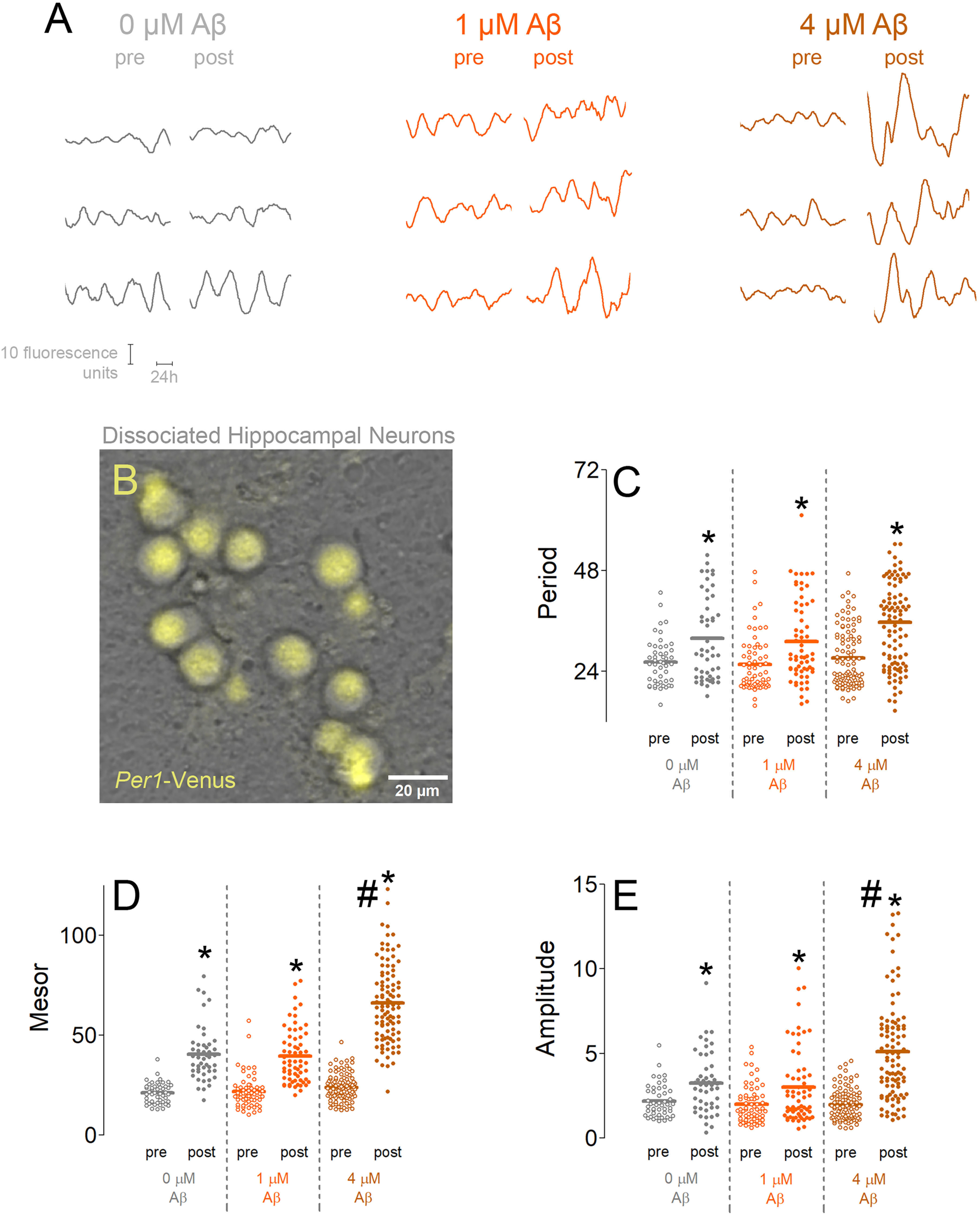
Administration of oligomerized Aß affects the expression and rhythm amplitude of *Per1*-Venus in cultured hippocampal neurons. Hippocampal neurons were isolated, plated and maintained in TTX-containing media. A: Representative Venus traces (n = 3 per treatment group) before (‘pre”) and during (“post”) treatment with either vehicle (0 µM) or oligomerized Aß (1 µM or 4 µM). B: Representative image of Venus-expressing hippocampal neurons; Venus expression (yellow fluorescence channel) was merged with the brightfield image. Quantitative analysis of *Per1*-Venus rhythm periodicity (in hours) (C), rhythm mesor (D) and rhythm amplitude (E) revealed that with time in culture there was significant period lengthening and increase in mesor and rhythm amplitude within a single hippocampal neuron. For statistical analysis, paired t tests were used to compare baseline (“pre”) and treated (“post”) values for each SCN neuron to determine the group mean effect of treatment (vehicle: 0 µM; Aß: 1 µM or 4 µM) for panels C, D, and E. ** denotes a significant difference between the corresponding “pre” and “post” values. To assess differences across the 0 µM, 1 µM or 4 µM treatments groups, the “post” value was subtracted from the “pre” value for each group, and then these differences were compared using the Student’s t test (0 µM vs. 1 µM or 4 µM; and 1 µM vs. 4 µM); # indicates changes that differed significantly from both the 0 µM and 1 µM Aß groups. Of note, the change in mesor (D) and amplitude (E) following 4 µM Aß treatment was significantly greater than the changes observed after 0 µM or 1 µM Aß treatment. A Bonferroni-adjusted significance threshold of p < 0.0083 was applied. Each data point represents a single SCN neuron (n = 48 for 0 µM; n = 63 for 1 µM; n = 99 for 4 µM Aß).

## 4. Discussion

The dysregulation of the circadian timing system is an early and pervasive feature of Alzheimer’s disease, manifesting as disrupted endocrine rhythms, sleep-wake cycle disturbances, increased daytime napping, and nocturnal restlessness, which significantly worsen patient quality of life (Musiek et al., 2015, 2018; Wu & Swaab, 2005). While these symptoms are well-documented, the underlying mechanisms by which AD pathology impairs the circadian timekeeping system have yet to be fully elucidated. Our study aimed to dissect this relationship by examining the specific effects of the Aβ peptide, a central player in AD pathogenesis, on the SCN central pacemaker and on a key peripheral forebrain oscillator population (hippocampal neurons). Our findings reveal a dichotomy: the SCN, both in vivo and ex vivo, demonstrates resilience to Aβ, while hippocampal neuronal oscillators exhibit dose-dependent vulnerability. These results suggest that the circadian disruptions characteristic of AD may not stem from a failure of the central clock itself, but rather from a pathological disruption/decoupling of peripheral brain clocks from the SCN, a conclusion with potential implications for understanding the cognitive and affective sequelae of the disease.

### 4.1 Aβ and the SCN clock

In the 5xFAD mouse model, we observed only a modest reduction in the free-running period in constant darkness compared to wild-type (WT) littermates. This short-tau phenotype is consistent with some previous studies using different Aβ transgenic mouse models-notably however, the reported effects of transgenic Aβ vary widely (Duncan et al., 2012; Song et al., 2015; Wisor et al., 2005). Interestingly, additional *in vivo* studies where the SCN was exposed to Aβ reported modest effects on clock-regulated physiology. Along these lines, in rats with grafted tissue that overexpress Aβ proximal to the SCN, a weakening in the amplitude of the activity and core body temperature rhythms was observed (Tate et al., 1992), while the microinjection of Aβ into the SCN of hamsters was reported to trigger increased variability in the phasing of locomotor activity onset, while no effects were observed on mesor, acrophase, or amplitude of the activity rhythm (Furio et al., 2002). Given these complex and somewhat contradictory sets of findings, we sought to add an additional set of *ex vivo* experiments that were designed to provide a crucial mechanistic layer to these behavioral observations. In these studies, which profiled SCN timing from 5xFAD tissue, as well as profiling via direct application of oligomerized Aβ, we did not detect a significant alteration in the endogenous period nor amplitude of molecular oscillations. Similarly, in dispersed SCN neurons, oligomerized Aβ did not affect clock periodicity or amplitude. Thus, in our model systems, the core timekeeping machinery of the SCN is intrinsically resistant to the physiological (and potentially pathological) actions of Aβ. Finally, it is worthwhile highlighting studies on Aβ plaque expression in the SCN of AD patients. In two studies, Aβ plaques were found in the SCN of AD patients, and a decrease in the neuronal cell density was reported (Harper et al., 2008; Stopa et al., 1999). In contrast, recent digital spatial profiling work indicates that Aβ plaques are absent from the SCN in AD patients throughout the progressive Braak stages (Son et al., 2024). Finally, while early stage AD patients are reported to have significantly disturbed sleep and activity patterns, their daily cortisol rhythm was found to be robust and unaffected (Hatfield et al., 2004). Given that both activity and cortisol rhythms are gated by the SCN clock, these data point to circuit-level dysregulation downstream of a resilient SCN that underlies dysregulated sleep and activity cycles in AD patients. Clearly, additional work both in animal models and human tissue will be required to create a clearer understanding of Aβ pathophysiology in the SCN.

### 4.2 Enhanced SCN Light Sensitivity

An interesting finding from our behavioral analysis was the enhanced sensitivity of 5xFAD mice to the entraining effects of light. These mice re-entrained to changes in the light cycle faster than their WT counterparts and could synchronize their activity to exceptionally dim light levels. This suggests that Aβ, while largely sparing the functional fidelity of the SCN clock, may sensitize the photic input pathway to the SCN. This pathway, originating in intrinsically photosensitive retinal ganglion cells (ipRGCs) that project to the SCN core, is the primary means by which the SCN is synchronized to the external light-dark cycle. Interestingly, a recent study by (Weigel et al., 2023) also reported heightened light sensitivity/entrainment of the clock in the 5xFAD mouse line, an effect that was associated with an increase in the density of ipRGCs. Of note, our light-evoked MAPK profiling assay (which we have used as a readout of light sensitivity/entrainment capacity of the SCN clock-Dziema et al., 2003; Butcher, et al., 2003) did not detect a heightened level of photic responsiveness in the SCN. The absence of an effect on the MAPK pathway may be due to our light stimulus paradigm, which might not have been properly calibrated in terms of intensity, duration, or circadian timing to reveal a phenotypic effect. Clearly, additional experiments that explore light-entrainment capacity using additional functional readouts (e.g., inducible gene expression, Ca^2+^ responsiveness) should help clarify the cellular and circuit processes that could contribute to this phenomenon.

Here, it is worth noting that, if heightened light sensitivity also occurs in AD patients, it could render the clock more susceptible to disruption by low levels of light at night. Consistent with this, increased light sensitivity could lead to the suppression of melatonin production and sleep disruption, thereby further contributing to circadian misalignments/dysregulation. As has been highlighted in a number of articles (Hanford & Figueiro, 2013; Zhu et al., 2022), carefully controlled light therapy, designed to provide a strong, appropriately timed signal while strictly avoiding light exposure at night, could be especially critical for AD patients.

### 4.3 Aβ and clock timing in the hippocampus

In contrast to our observations of resilient SCN timing, hippocampal cellular oscillators exhibited a vulnerability to Aβ exposure. Specifically, ex vivo application of Aβ induced a dose-dependent modulation of circadian oscillations, which manifested as higher amplitude rhythms and an elevated baseline level of the *period1* reporter.

Numerous studies have demonstrated that critical functional properties of forebrain neurons are regulated by the circadian clock. For example, circadian phase modulates long-term potentiation (LTP) induction thresholds, membrane capacitance, and plasticity-associated signaling pathways (Chaudhury et al., 2005; Eckel-Mahan et al., 2008; Hoyt & Obrietan, 2022; Munn et al., 2015; Nakatsuka & Natsume, 2014; Severin et al., 2024; Shimizu et al., 2016). At the behavioral level, disruption of hippocampal clock function has profound consequences; Targeted deletion of *Bmal1* in excitatory cortical and limbic neurons, while sparing SCN function, results in marked impairments in long term memory (Price, & Obrietan, 2018; Shimizu et al., 2016).

At the level of cellular timing, it is not clear why GABAergic SCN neurons exhibit resistance, whereas hippocampal neurons are sensitive to the effects of Aβ. Potential explanations could be related to the differential expression of receptors and signaling pathways that mediate Aβ toxicity, such as NMDA receptors and/or pathways involving cellular prion protein and Ca^2+^ homeostasis; Reduced expression could render the SCN less susceptible to Aβ-induced stress and Ca^2+^ overload than hippocampal neurons. Relatedly, in the hippocampus, Aβ can aberrantly increase extracellular glutamate levels and drive excessive NMDA receptor activation and Ca^2+^ dysregulation (Alberdi et al., 2010; Danysz & Parsons, 2012; Mattson et al., 1992), which would affect a range of downstream intracellular signaling pathways. Of note, the expression of the *period* class of genes (as well as other core clock genes) is regulated by Ca^2+^-sensitive CaMK and MAPK signaling (Akashi & Nishida, 2000; Kon et al., 2014; Nomura et al., 2003; Obrietan et al., 1998). As such, one could envision a signaling pathway by which Aβ-evoked cell stress could lead to increased expression of the *period1* gene. The resulting effects of elevated levels of Period1 and higher amplitude clock oscillations are likely to be multifaceted. For example, perturbation of clock rhythms in the hippocampus (e.g., increased amplitude and mesor) would likely dysregulate the expression of clock-controlled genes, which in turn could impair circadian modulation of neuronal excitability and plasticity-two processes that are fundamental to cognition. Collectively, these data suggest that beyond its established neurotoxic effects, Aβ also undermines fundamental circadian regulatory processes within hippocampal circuits. Further, at the systems level, Aβ-induced disruption could decouple corticolimbic oscillators from SCN-derived entrainment cues, leading to broader network desynchrony (discussed further below). This dual impact on cellular viability and temporal coordination may represent a critical and underappreciated mechanism contributing to cognitive decline in AD pathogenesis.

### 4.4 Decoupling of Circadian Circuits: A Systems-Level View of Cognitive Decline

The key implication of our research is the concept of Aβ-mediated "decoupling" of circadian circuits. We propose a model where the SCN continues to generate a robust, albeit slightly altered, timing signal, but this signal fails to effectively entrain downstream oscillators in corticolimbic circuits that have been dysregulated (and in turn, uncoupled) by Aβ. The result is a brain operating in a state of internal desynchrony. The hippocampus, for instance, can no longer align its memory-encoding machinery with the optimal circadian phase dictated by the SCN. This temporal mismatch could directly contribute to the learning and memory deficits characteristic of AD. For example, if the SCN signals that it is the appropriate time for memory consolidation (i.e., during sleep), but the hippocampal clock is incapable of receiving or acting on this cue, the consolidation process would be severely impaired. This decoupling framework extends beyond the hippocampus to other cognitive domains. Peripheral oscillators exist in numerous brain regions, including the prefrontal cortex and amygdala, which govern executive function and emotional regulation, respectively (Albrecht & Stork, 2017; Liu et al., 2024; Puig et al., 2023). The disruption of their intrinsic clocks by Aβ would lead to a similar functional disconnection from the SCN, potentially providing insights into the impairments in executive control and the affective disturbances (e.g., anxiety, depression, apathy) that are also prominent in AD. From this perspective, circadian disruption is not merely a symptom of AD but an integral part of its pathophysiology, which manifests as a systems-level failure of temporal organization that could compromise the system and circuit level processes across the CNS. Additional work that supports these models would potentially reframe our understanding of AD-related cognitive decline, suggesting that restoring the synchronized timing of neural circuits, perhaps by targeting the health of peripheral oscillators or by amplifying the SCN output signal, could represent a novel therapeutic strategy to ameliorate the cognitive and behavioral symptoms of the disease.

In conclusion, our findings raise the prospects that cellular and circuit-level disruption underlies circadian disturbances seen in AD. By demonstrating the resilience of the central SCN clock and the vulnerability of hippocampal oscillators to Aβ, we shift the focus from a "broken" SCN clock to a model centered on system and circuit level ‘decoupling’. This work supports the importance of investigating not just the core pacemaker but the entire distributed network of brain clocks to understand, and ultimately treat, the temporal disorganization that is a key feature of Alzheimer’s disease.

## Acknowledgements

Anisha Kalidindi and Yen Anh Nguyen, for assistance with animal management and with the preparation of the dot-blot.

## 5. Author Contributions

**Kari R. Hoyt:** Data curation, Formal analysis, Funding acquisition, Investigation, Methodology, Project administration, Supervision, Visualization, Writing – original draft, Writing – review and editing. **Tyler Kyhl:** Data curation, Investigation. **Nicklaus R. Halloy:** Data curation. **Karl Obrietan:** Data curation, Formal analysis, Funding acquisition, Investigation, Methodology, Project administration, Supervision, Visualization, Writing – original draft, Writing – review and editing.

## 6 Glossary

5xFAD mouse line: (A transgenic mouse model of amyloid pathology)
Aβ: Amyloid Beta
AD: Alzheimer’s Disease
Bmal1: (Protein that functions as a key component of the core molecular circadian clock)
Clock: (Protein that functions as a key component of the core molecular circadian clock)
CRY: Cryptochrome (Protein that functions as a key component of the core molecular circadian clock)
DAB: Diaminobenzidine
ELISA: Enzyme-Linked Immunosorbent Assay
ERK: (A kinase in the MAPK pathway)
GFP: Green Fluorescent Protein
ipRGCs: intrinsically photosensitive retinal ganglion cells
LL: (Constant light)
Lux: (Unit of light measurement)
MAPK: Mitogen Activated Protein Kinase (a cell signaling pathway)
PBS: Phosphate Buffered Saline
PER: (protein that functions as a key component of the core molecular circadian clock)
*Per1*-Venus: (Transgene used in our circadian reporter mouse model)
SCN: Suprachiasmatic Nucleus
DD: (Total darkness)
YFP: Yellow Fluorescent Protein

